# Structural Insights into Zn^2+^ and Ca^2+^ binding to Human Protein Z and Their Impact on Membrane Association

**DOI:** 10.1101/2025.10.09.681514

**Authors:** Subash Chellam Gayathri, Suchetana Gupta, Tanusree Sengupta, Soumyadev Sarkar

## Abstract

Calcium ions (Ca²⁺) bound to the γ-carboxy glutamic acid-rich (Gla) domain of coagulation proteins are essential for their membrane anchoring. Zinc ions (Zn²⁺), abundant in plasma, also bind coagulation proteins and modulate their activity. However, their role in membrane-protein interactions remains less understood. The current study examines how Ca²⁺ and Zn²⁺ influence the structure and function of human protein Z (PZ), a cofactor of protein Z–dependent protease inhibitor (ZPI) that inhibits factor Xa on phospholipid surfaces. Zn^2+^ was found to substitute for Ca^2+^ in the Gla domain of PZ and promote its membrane binding by inducing conformational changes like Ca^2+^. Additionally, a Zn^2+^ located in the epidermal growth factor1 (EGF1) domain of PZ appeared to synergize with Ca^2+^ to enhance the membrane affinity of PZ. Furthermore, both Ca^2+^ and Zn^2+^ were suggested to destabilize PZ-ZPI complex, thereby modulating the cofactor activity of PZ, a finding that warrants further experimental investigations. Collectively, this study provides the first structural and functional characterization of Zn²⁺-bound PZ, offering new insights into metal ion driven structural dynamics of PZ and their implications in coagulation regulation.

## 1. Introduction

Blood coagulation involves a series of enzymatic reactions that occur on the membrane surfaces. Activated membranes accelerate these reactions upon binding and aligning various clotting factors, cofactors and anticoagulants congruously on their surface. Calcium (ions) play a pivotal role in the protein-membrane interaction of nearly all vitamin K dependent (VKD) coagulation proteins. Primarily, Ca^2+^ facilitates the proper folding of the γ carboxy glutamic acid rich (Gla) domain which is essential for the membrane association of VKD proteins. Additionally, Ca^2+^ induces anionic phospholipids, especially phosphatidyl serine (PS) molecules, to expose negatively charged areas on activated phospholipid membranes. This exposure enables calcium’s bivalent charge to anchor the Gla domain firmly to the membrane. While internal calcium ions stabilize the optimal structure of the Gla domain for membrane binding, external calcium ions are directly involved in the membrane attachment process. Most studies have used Ca^2+^ concentrations ranging from 2 mM to 5mM for the assembly and formation of different coagulation complexes despite the physiological plasma concentration of Ca²⁺ being approximately 1 mM [1]. However, plasma also contains about 0.6 mM magnesium ions (Mg²⁺), which occupy specific binding sites within the Gla domain and support the binding of clotting factors to phospholipid membranes. Thus, at suboptimal calcium concentrations, Mg²⁺ plays a crucial role in enhancing the affinity of coagulation factors VIIa (fVIIa), IXa (fIXa), and prothrombin for lipid membranes [2].

Ca^2+^ binding sites have also been identified in other structural domains of most of the VKD proteins. These high affinity Ca^2+^ binding sites often enhance the protein’s affinity for their cofactors, thereby regulating their activity. Both fVIIa and fIXa possess high affinity Ca^2+^ binding sites in both the protease domain and the EGF1 domain [3]. A high affinity Ca binding site located outside the Gla domain of Protein C is absolutely essential for its conversion to the activated form (APC) [4]. In most cases, these high affinity Ca^2+^ ions cannot be replaced by other metal ions. However, coagulation proteins may also have potential binding sites for other bivalent metal ions, and binding of these ions often alters their activity. For example, Zn impairs the inhibitory activity of APC towards both fVa and a synthetic substrate, probably, by binding to a site in protease domain distinct from the Ca^2+^ binding site [5]. Zn^2+^ has been reported to abate the amidolytic activity of fVIIa by binding to its protease domain [6]. In the presence of Zn^2+^, Protein S efficiently binds to fXa and inhibits thrombin generation by fXa/fVa (prothrombinase complex) [7]. Additionally, Zn^2+^ enhances fXII autoactivation by inducing a conformational change in the protein [8]. Zn^2+^ also induces a conformational change in the D5 domain of high molecular weight kininogen (HMK), another protein involved in the contact pathway of blood coagulation, likely facilitating HMK binding to polyanionic surfaces [9].

Human Protein Z (PZ) is a serine protease inhibitor homolog whose domain structure closely resembles that of other VKD proteins such as factors VIIa, IXa, Xa, featuring a Gla domain and two epidermal growth factor like domains (EGF1 and EGF2). However, PZ cannot function as a serine protease inhibitor as the His and Ser residues of the catalytic triad, which are present in other coagulation proteases, are missing in the homologous region of PZ [10]. Instead, PZ acts as a cofactor of a serine protease inhibitor known as protein Z dependent protease inhibitor (ZPI), which inhibits membrane bound fXa [11]. A nearly thousand-fold enhancement of fXa inhibition by ZPI is observed in the presence of PZ, procoagulant lipid and Ca^2+^. During membrane activation, the PZ-ZPI complex binds to the membrane surface through the Gla domain of PZ, simultaneously co-localizing ZPI near the membrane [12]. This interaction increases the local concentration of ZPI near the membrane surface and positions it close to fXa, which is bound to the same membrane as PZ. It has been previously demonstrated that PZ binds to two pivotal membrane phospholipids, phosphatidyl serine (PS) and phosphatidyl ethanol amine (PE), with equal affinity in a Ca^2+^ dependent manner [13]. Evidently, Ca^2+^ ions bound to the PZ-Gla domain, are critical for its membrane binding and subsequent cofactor activity. Notably, bovine PZ lacking Gla domain does not possess any Ca^2+^ binding sites, as reported by Morita et al [14. Therefore, it can be assumed that no Ca^2+^ binding sites exist in PZ apart from those in its Gla domain. However, whether other bivalent metal ions such as Zn^2+^, can bind to PZ and alter its function, as they do for some other coagulation proteins, remains to be explored.

The current study, therefore, investigates the binding of Zn^2+^ to PZ and the structural changes induced thereby. Both computational and experimental approaches were employed to visualize Zn^2+^ binding in detail. We demonstrate that Zn^2+^ binds to PZ with a dissociation constant (k_d_) of 122 ± 12 µM, and this binding exhibit synergy with Ca^2+^. Our experimental data further suggest that soluble lipids can bind to PZ in the presence of Zn²⁺ even without Ca²⁺. We also examined the binding of Mg^2+^ to PZ, however, only very weak binding was observed and so we did not pursue further studies with Mg^2+^.

Next, we constructed the full-length structural model of Zn²⁺ and Ca^2+^-bound PZ which were subjected to molecular dynamic (MD) simulations. Computational analyses identified Zn^2+^ binding sites in both Gla and EGF1 domain of PZ. The ionic interactions involving Zn²⁺ in the EGF1 domain prevented PZ from adopting the fully folded conformation seen in the presence of Ca²⁺. Additionally, Zn²⁺ binding in the EGF1 domain exposed the keel region towards the exterior, potentially enhancing PZ’s interaction with the lipid membrane. Zn²⁺ binding in the EGF1 domain may be functionally significant, as it could disrupt the stabilizing interaction between PZ and ZPI. Collectively, the structural insights gained from this study provide a solid foundation for understanding how the bivalent metal ions, Ca²⁺ and Zn²⁺, regulate the structural conformation and membrane binding of PZ, either independently or synergistically. This understanding is crucial for elucidating the mechanism of PZ-ZPI complex formation on phospholipid membranes.

## 2. Materials and Methods

### 2.1. Materials

Human PZ was purchased from r2 diagnostics (sister company of Enzyme Research Laboratories, IN, USA). The sodium salt of 1, 2-dicaproyl-sn-glycero-3-phospho-L-serine (C6PS) was purchased from Avanti Polar Lipids (Alabaster, AL, USA). All other chemicals were from sigma chemicals and of best available grade.

### 2.2. Fluorescence Measurements

Intrinsic tryptophan fluorescence of PZ in appropriate buffer (20 mM tris-HCl, 150 mM NaCl, 0.6% PEG, pH7.4) was measured as a function of increasing concentration of Zn^2+^ in absence and presence of Ca^2+^ (2 mM) using a FluoroLog spectrofluorometer (Horiba Jobin-Yvon Inc., Edison, NJ) with an excitation wavelength of 285nm (band-pass 4 nm) as described in [15] Emission spectra were recorded at 340 nm (band-pass 4 nm). PZ incubated with three different concentrations of Zn^2+^ (0.5 mM, 1mM, and 3mM) were again titrated with varying concentrations of soluble C6PS using the same method as described above. Lipid titrations were measured below the critical micelle concentration (CMC) of C6PS. Apparent dissociation constants for binding of PZ to metal ions and/or soluble lipids were obtained by fitting the experimental data to a simple, single-site binding model.

### 2.3. Model building

In order to model the full-length protein, we originally employed AlphaFold server [16]. However, the resulting structures could not predict any folded conformation of the Gla domain and were considerably entangled. Therefore, we resorted to model building using SWISS-MODEL as described. An initial template search for the Gla domain of PZ was performed using the primary sequence of the Gla domain with BLAST and HHblits in the SWISS-MODEL terminal. A homologous structure of a folded Gla domain in human coagulation fVIIa (PDB ID 2A2Q) was chosen as the template [17]. Primary sequence identity between the PZ and the template was 49 % with a query coverage of 94 %. The generated model of the Gla domain of PZ was tested for acceptable quality. The model was then linked to the EGF1 domain in the crystal structure of PZ (PDB ID 3H5C) using Coot [18]. The final model was subject to geometry minimization in Phenix [19] prior to MD simulation using AMBER package for solvation, and for additional all atom minimization to remove any poor/overlapping contacts.

### 2.4. Molecular Dynamic Simulations

MD simulations were conducted on the generated systems using AMBER 18 [20] with the AMBERff14 SB force field [21]. Initially, all structures underwent energy minimization for 2000 steps using the steepest descent and conjugate gradient algorithms. The minimized structures were then solvated in a cubic water box containing TIP3P water molecules [22]. To neutralize the system charges, an appropriate number of Na^+^ and Cl⁻ ions were added. Electrostatic interactions were calculated with a 12 Å cutoff using the Particle Mesh Ewald method [23].

Preliminary minimization steps were performed to eliminate unfavorable contacts, followed by equilibration in the NVT ensemble at 300 K for approximately 500 ps. Subsequently, system density was equilibrated under NPT conditions at 1 atm pressure for 1 ns. A time step of 2 fs was maintained throughout the minimization, equilibration, and production phases. Once the energy and density values stabilized, the systems were subjected to a 100 ns production run in the NPT ensemble at 300 K and 1 atm pressure. Coordinates were recorded every 2 ps. Trajectory visualization was carried out using VMD [24], while analyses including RMSD, RMSF were performed using the CPPTRAJ module of AMBER12 [25].

## 3. Results and Discussion

### 3.1. Binding of Zn^2+^ to PZ

The interaction between PZ and Zn²⁺ was examined by monitoring changes in the intrinsic tryptophan fluorescence intensity of PZ as a function of increasing Zn²⁺ concentration, with the excitation wavelength fixed at 295 nm. The relative fluorescence intensity was plotted against Zn²⁺ concentration (●), as shown in Fig. 1. A progressive decrease in fluorescence intensity was observed with rising Zn²⁺ levels, suggesting that Zn²⁺ binding induces conformational changes in PZ that affect the local environment of its tryptophan residues. These conformational alterations likely reflect structural rearrangements upon Zn^2+^ coordination. The fluorescence data fitted to a single-site binding model, yielded a dissociation constant (K_d_) of 122 ± 12 μM, indicative of a moderate affinity of Zn²⁺ for PZ under these conditions.

**Fig. 1.**
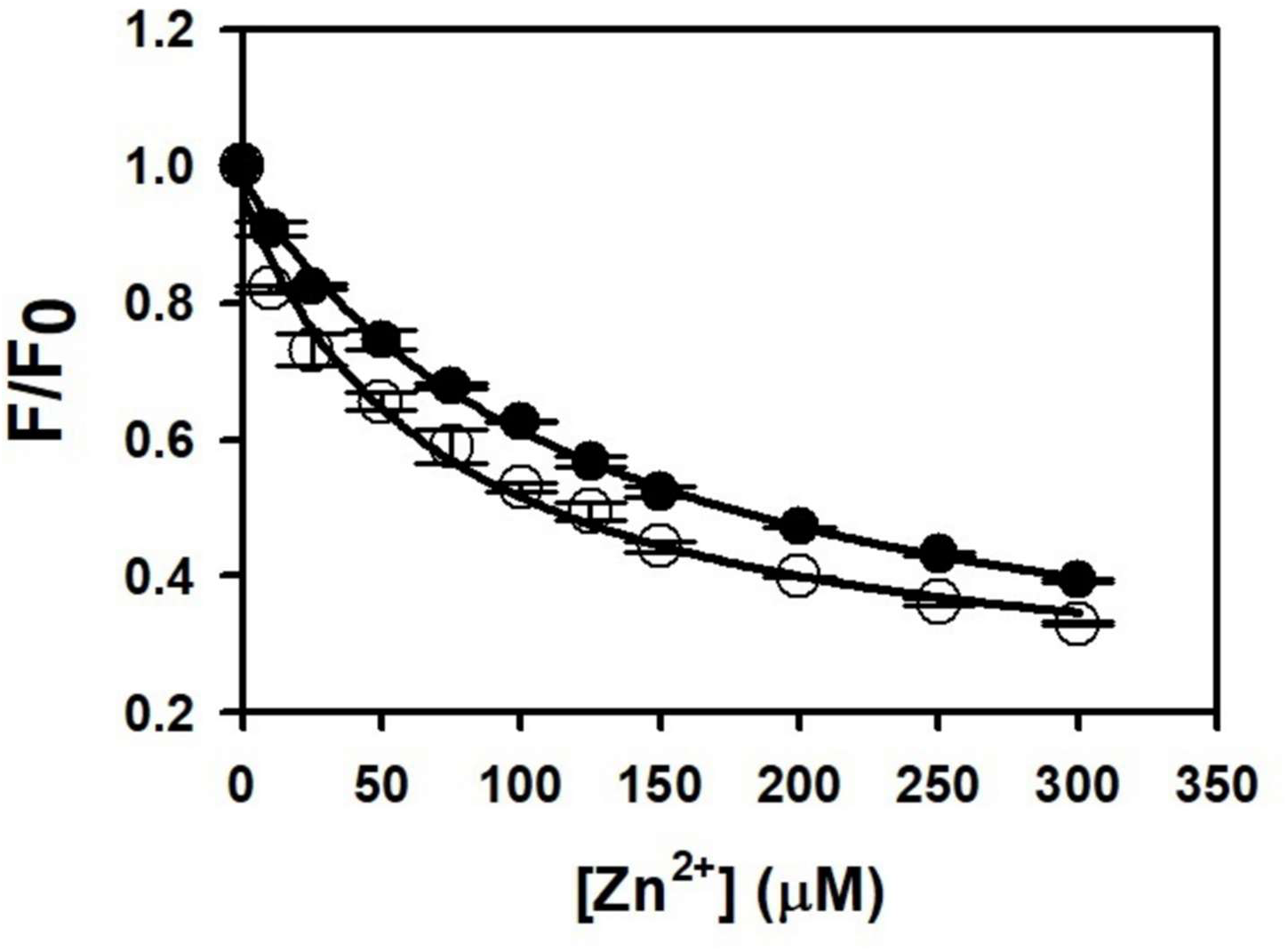
Binding of Zn^2+^ in absence (●) and presence of Ca^2+^ (○) to PZ as detected by intrinsic fluorescence. Integrated intrinsic fluorescence of 100 nM PZ in 20 mM tris-HCl, 150 mM NaCl, 0.6% PEG, pH 7.4 was measured as a function of Zn^2+^ (●) concentration to monitor binding. The single site binding model provided an apparent K_d_ of 122 ± 12 µM. The same experiment was repeated by titrating PZ preincubated with 2 mM Ca^2+^ with increasing concentration of Zn^2+^ (○). Single site analysis yielded an apparent K_d_ of 70 ± 13 µM indicating synergistic binding of Zn^2+^ in presence of Ca^2+^. Each data point is mean ± SD of three replicates.

To evaluate the effect of Ca²⁺ on Zn²⁺ binding, PZ was preincubated with 2 mM Ca²⁺ for 15 minutes prior to titration with Zn²⁺ under identical conditions. The fluorescence intensity changes were recorded and analyzed similarly, with the resulting binding curve shown in Fig. 1 (datapoint shown in ○). Fitting this data to the same single-site binding model produced an apparent K_d_ of 70 ± 13 μM, which is significantly lower than the K_d_ observed in the absence of Ca²⁺. Enhancement of the affinity of PZ for Zn²⁺ in presence of Ca²⁺ suggests a synergistic effect between these two divalent metal ions in modulating PZ structure and function.

### 3.2. Binding of soluble lipids to PZ in presence of Zn^2+^

Lipid binding by coagulation proteins typically requires the presence of Ca^2+^. However, under conditions of limited Ca^2+^ availability, other bivalent cations such as Mg^2+^ have been shown to facilitate this interaction [26]. Given the demonstrated affinity of Zn²⁺ for PZ, we investigated whether Zn²⁺ could similarly support the lipid-binding activity of PZ. The effect of Zn²⁺ on lipid binding by PZ was assessed using soluble lipids, which are widely employed as effective membrane mimetics for studying lipid-protein interactions in coagulation. Although these soluble lipids do not occur physiologically, they provide a valuable tool for monitoring events that are challenging to study in membrane systems. Previous studies have successfully utilized soluble lipids to identify and characterize lipid-binding sites on several coagulation proteins, including fXa, fIXa, fVa, fVIIa [15,27]. For the current experiments, only C6PS was used, given the comparable binding affinities of PZ for C6PS and C6PE [13]. Fluorescence emission spectra of PZ were recorded as a function of increasing C6PS concentration in buffer (20 mM tris-HCl, 150 mM NaCl, 0.6% PEG) lacking Ca^2+^ but containing varying concentrations of Zn^2+^ (500 μM, 1mM, and 3 mM). The K_d_ values for C6PS binding to PZ were determined to be 160, 147, 94 μM in the presence of 500 μM, 1mM, and 3 mM, respectively (Table 1). These results demonstrate that, in the absence of Ca²⁺, Zn²⁺ can significantly enhance the binding of PZ to lipid membranes, suggesting a potential role of Zn²⁺ in modulating PZ-lipid interactions under physiological or pathophysiological conditions.

**Table 1.** Soluble Phosphatidylserine (C6PS) binding to PZ in presence of different concentrations of Zn^2+^. The data are mean± SD of three replicates.

| Titrant | $\text{Zn}^{2+}$ (mM) | $K_d$ ($\mu\text{M}$) |
| --- | --- | --- |
| C6PS | 0.5 | $160 \pm 23$ |
| C6PS | 1.0 | $147 \pm 17$ |
| C6PS | 3.0 | $94 \pm 12$ |

### 3.3. Model building

Previous structural and functional studies of VKD coagulation proteins revealed that the structural rearrangement of the Gla domain induced by Ca²⁺ ions facilitate membrane anchoring, maintains the domain’s optimal folded conformation in solution [17]. Our experimental studies further demonstrate that Ca^2+^ and Zn^2+^ act synergistically to amplify the membrane binding of PZ. Thus, we propose that metal ion binding induces conformational changes in the PZ’s Gla domain, aiding membrane binding and functional enhancement thereof. However, the experimental structures of PZ solved to-date lack the Gla domain (PDB ID: 3F1S and 3H5C), limiting molecular insights into these conformational shifts. Consequently, to validate our experimental data and to explore the conformational landscape accessed by PZ, we resorted to MD simulations on a hypothetical model of full-length PZ structure in the presence of Ca^2+^ and Zn^2+^ independently and together. This approach enables atomic-level characterization of metal ion-induced structural changes previously inaccessible through experimental methods.

PZ, like other homologous VKD proteins, contains a serine protease (SP) like domain that is pseudo-catalytic due to the absence of the obligatory catalytic triad residues, two EGF domains along with the intrinsically flexible Gla domain (Fig. 2A) [28]. Homology modeling identified Gla domain of coagulation factor VIIa (PDB ID: 2A2Q) as the best template for PZ-Gla domain. The resulting Gla domain model was evaluated for quality and connected to the coordinates of EGF1 domain obtained from the PZ crystal structure. The final full length Gla domain incorporated structural model did not perturb the EGF and SP domains of PZ and retained all the disulfide bridges that were present in the original experimental structures. The last frame of the model was used for subsequent MD runs with the metal ions for functional predictions. The simulated structures were found to be quite stable as observed from the root mean square deviation (RMSD) plot (Fig. S1).

**Fig. 2.**
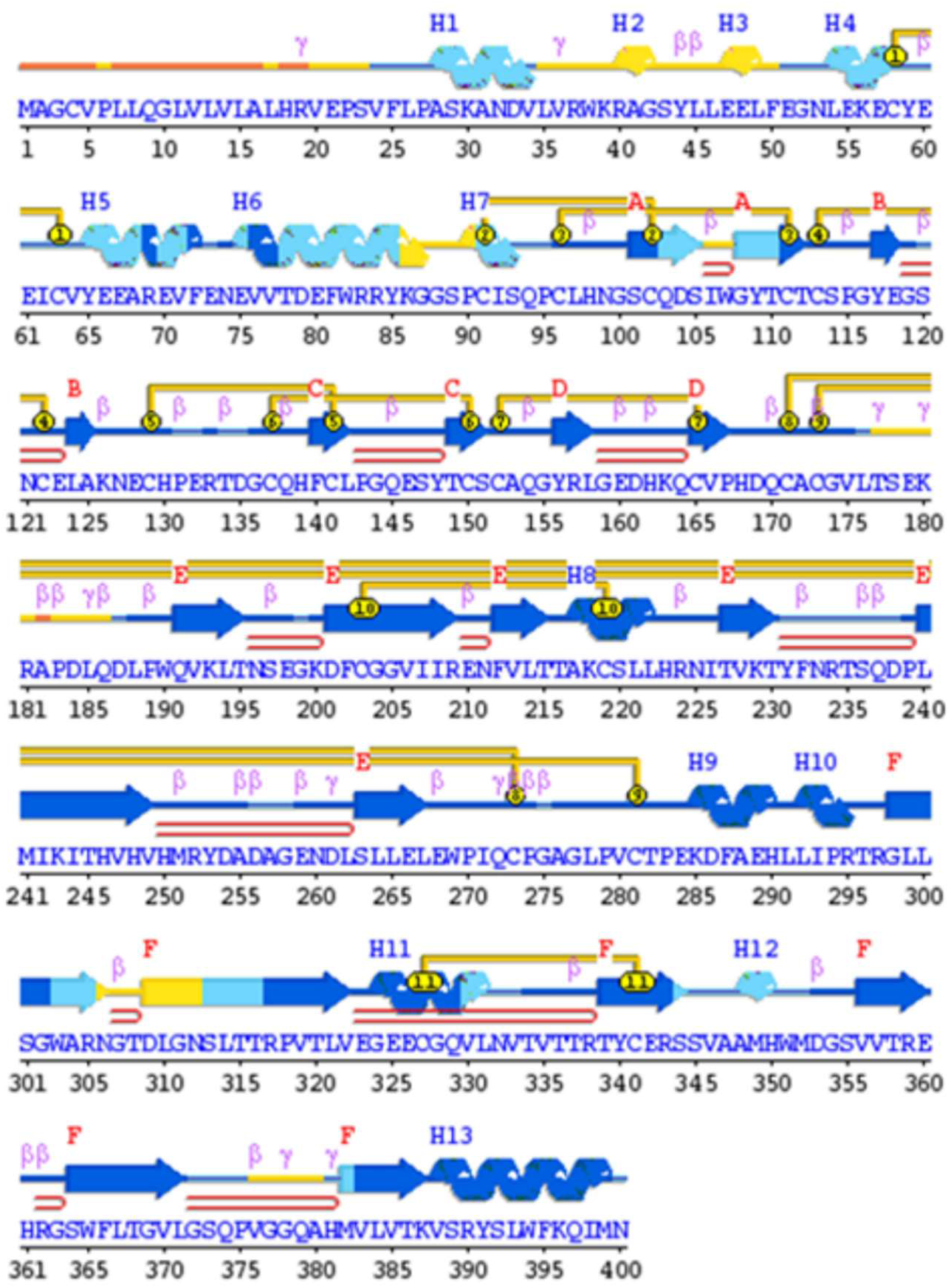
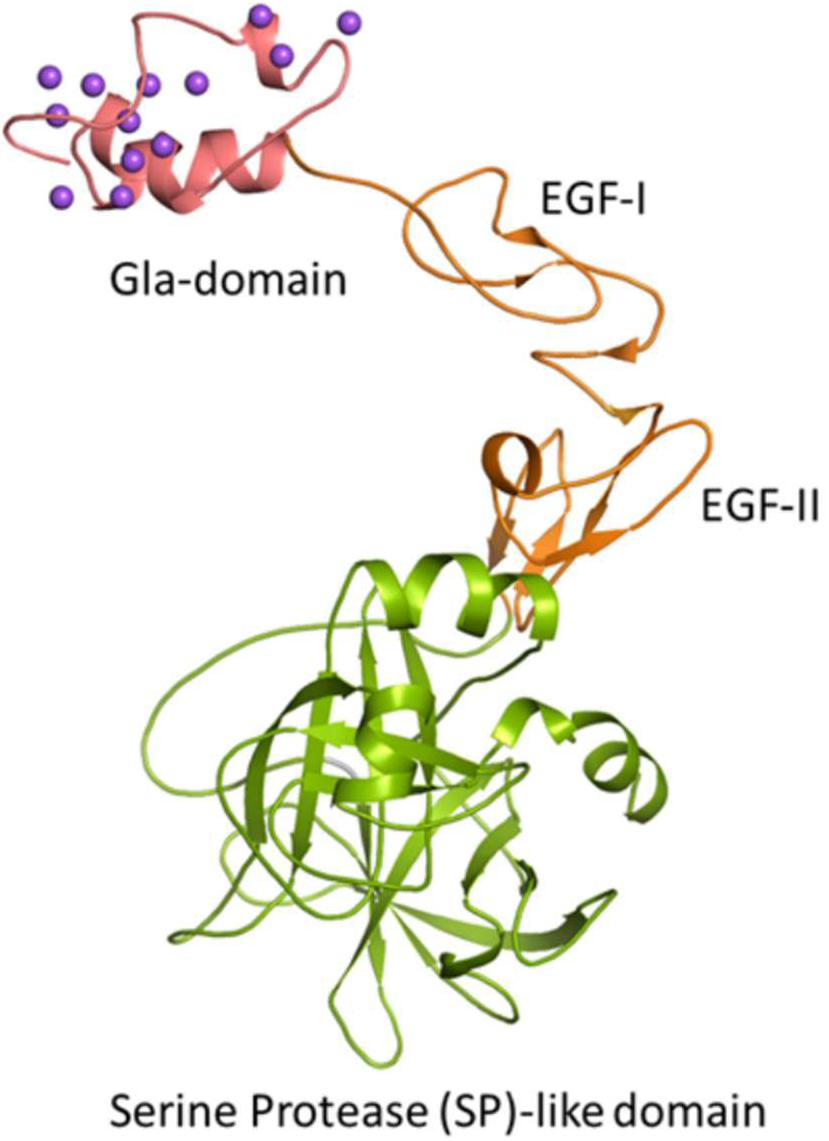
Structure of Human Protein Z (PZ). A. Representation of primary and secondary structure of full-length PZ (UniProt ID: P22891). Residue numbers corresponding to different domains are as follows: Gla domain (41-86), EGFI (87-123) and EGF II (125-166), SP-like domain (175-400). Disulphides are shown as yellow connected lines. Secondary structures, alpha helices (H) and beta sheets (arrows, ᵝ, ˠ) are shown above the residues. Figure generated in PDBsum using the AlphaFold model of ProZ as template. Alphafold confidence scores are shown as variable colours of the secondary structure – dark blue (very high) light blue (confident) yellow (low) and orange (very low). **B. Cartoon representation of the MD simulated, stable, full length structural model of PZ.** Gla domain (1-46) is shown in pink, EGFI (47-83) and EGF II (85-126) are in orange, SP-like domain (135-360) is in green, and 13 Na^+^ ions are shown as purple sphere (numbering is according to our structural model). Na^+^ ions occupying the Gla domain help maintain its secondary structure, however, Gla domain along with the EGF domain are bit extended and positioned away from the SP-like domain.

### 3.4. MD-Simulation studies

#### 3.4.1. Ca^2+^ binding to PZ

After a 100 ns MD simulation run, the full-length PZ model was found to pick 13 Na^+^ ions from the uniformly distributed Na^+^ ions of the solvent to stabilize the 13 gamma glutamate residues present in the Gla domain as evident from the Fig. 2B. This interaction of the Na^+^ ions assists in the retention of the folded conformation of the Gla domain that is usually observed in the presence of Ca^2+^ in homologous proteins [29]. Therefore, the coordinates of the Na^+^ ions were used to guide the docking of Ca^2+^ ions, and the Ca^2+^ loaded protein was again subjected to energy minimization in Phenix to allow the carboxy glutamate residues to adopt optimum geometry required for Ca^2+^ coordination. This initial energy minimized structure was then subjected to another set of MD run to investigate any conformational changes triggered by calcium binding.

In the stabilized output of the MD run, 11 Ca^2+^ ions were observed to be present in the Gla domain (Fig. 3). This is more compared to 6-7 Ca^2+^ ions that were observed for other VKD coagulation proteins [2, 26]. We observed a remarkable conformational change in PZ upon binding of Ca^2+^ ions to the Gla domain, compared to the Na^+^ bound structure. In the Na^+^ bound apo protein, both Gla and EGF domain were extended and thus positioned further away from the SP like domain. However, a large domain rotation of ∼130° at the N-terminus of the EGF1 domain (residues GGSP, 47-50) was observed in the Ca^2+^ bound structure, resulting in the reorientation of the Gla domain adjacent to the SP like domain of PZ (Fig. 3). Residues, Glu283 and Glu289, from the SP domain also contributed to the calcium coordination and perhaps stabilized the folded conformation of PZ (Fig. S2). Another interesting feature of the Ca^2+^ loaded model is that the probable “keel” residues Leu9 and Phe10 are exposed to the solvent and along the face of these residues are also a couple of solvent exposed Ca^2+^ ions coordinated by CGU15, CGU20 and CGU21 (Fig S2). Protruded keel residues and solvent exposed Ca^2+^ are considered to be important features for membrane insertion in homologous proteins and therefore it is likely that this face of the Gla domain is the most likely membrane interacting face of PZ [30].

**Fig. 3.**
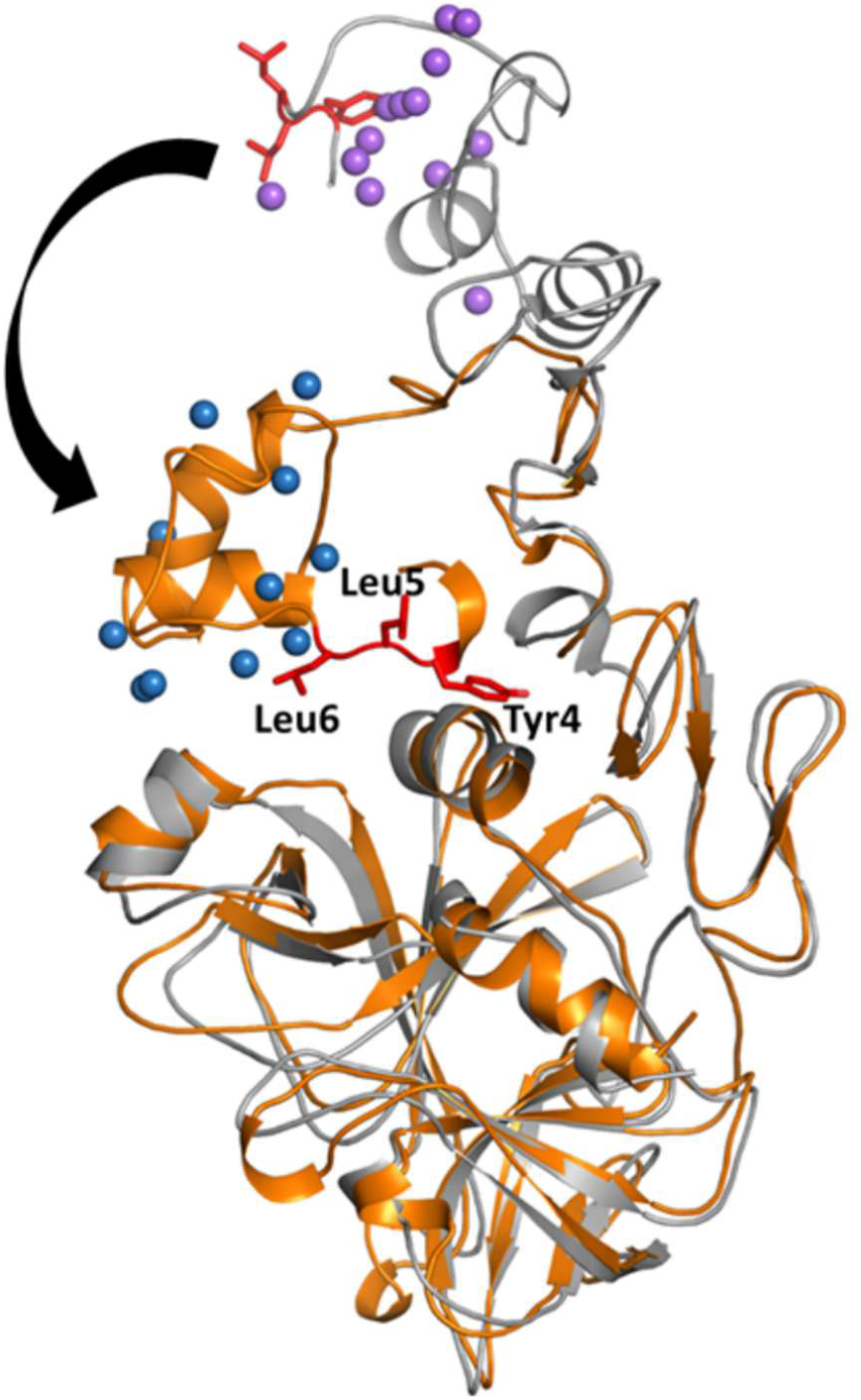
Reorientation of Gla domain in presence of Ca^2+^. A large domain rotation of EGFI was observed when Na^+^ ions (purple sphere) are replaced with Ca^2+^ ions (blue spheres) and subjected to MD simulation, leading to a repositioning of the Gla domain near the SP-like domain. 11 Ca^2+^ ions were seen to occupy the Gla domain among which the solvent exposed ones are believed to be involved in membrane insertion. Ca^2+^ bound PZ structure is shown in orange whereas Na^+^ bound in grey.

The N-terminal residues Tyr4, Leu5, Leu6, of the Gla domain are oriented towards the SP domain as observed in Fig. 3. Tyr4, together with His250 and His90 of the SP domain create a pocket (Fig. 4). Intriguingly, it is near this pocket that the ZPI interface “hot spot” residue Tyr240 interacts with PZ. Literature reports indicate that any changes in the interactions at this interface residue can influence RCL dynamics [31]. We performed two additional MD runs on a modelled PZ-ZPI complex, using the coordinates of ZPI from the experimental crystal structure (PDB ID: 3F1S) in complex with the Ca^2+^ loaded compact form and Ca^2+^ free extended form of PZ. Interestingly, the overall root mean square fluctuations (RMSF) of the closed compact form in complex with ZPI was higher in comparison to the open extended form (data not shown), indicating reduced stability of PZ-ZPI complex in presence of Ca^2+^. As the Ca^2+^ bound closed compact form of PZ would bind more tightly to the membrane, it may exhibit lower affinity for ZPI. Consequently, this could enhance the ability of ZPI to interact with and inhibit its substrate protease, fXa, when localized on the same membrane surface as PZ. The physiological significance of these findings, particularly the role of Ca²⁺ binding to PZ in the assembly of the PZ–ZPI complex, requires further experimental investigation in the future.

**Fig. 4.**
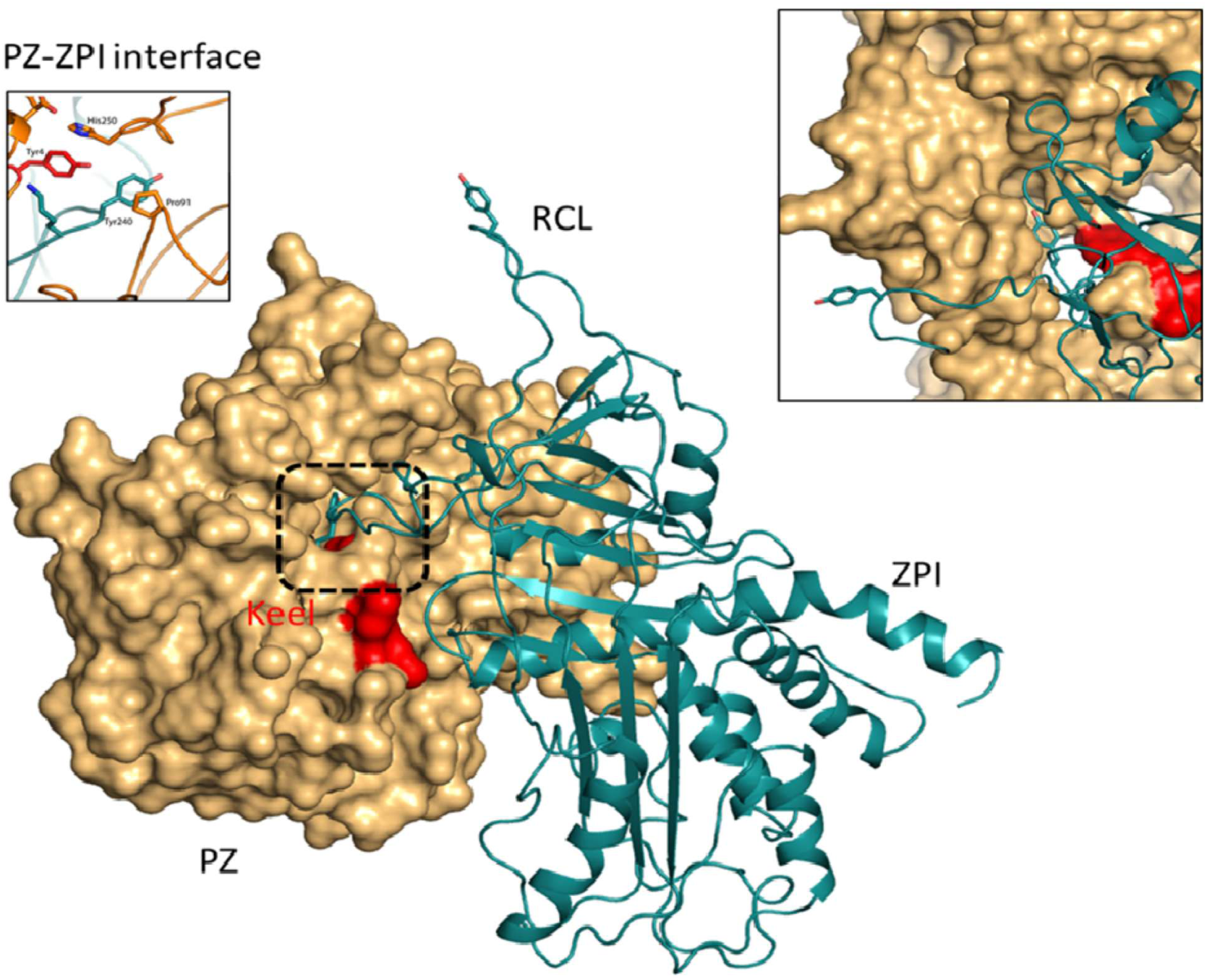
Structure of the complex between ZPI and Ca^2+^ bound PZ. PZ is shown as a beige surface representation with the keel region highlighted in red. ZPI is shown as blue ribbons. The ZPI hotspot Tyr240 residue is observed to bind near a pocket with residues from the keel region. Inset shows the PZ-ZPI interface residues.

#### 3.4.2. Zn^2+^ binding to PZ

To elucidate the binding of Zn^2+^, MD-simulations were performed under various conditions. First, we replaced the Ca^2+^ ions by Zn^2+^ in the Ca^2+^ loaded, closed PZ, and the structure was subjected to MD simulation. Zn^2+^ was observed to occupy the Ca^2+^ binding sites with minimal structural changes (Fig. 5), which aligned well with our experimental findings showing similar binding constants for both Zn^2+^ and Ca^2+^. Furthermore, analysis of the metal ion bound structures indicated that once Ca^2+^ binding induces the folded conformation, Zn^2+^ can subsequently replace Ca^2+^ ions depending on spatiotemporal context. These structural features confirmed the synergistic binding of the two ions, as predicted by our tryptophan fluorescence studies (Fig. 1).

**Fig. 5.**
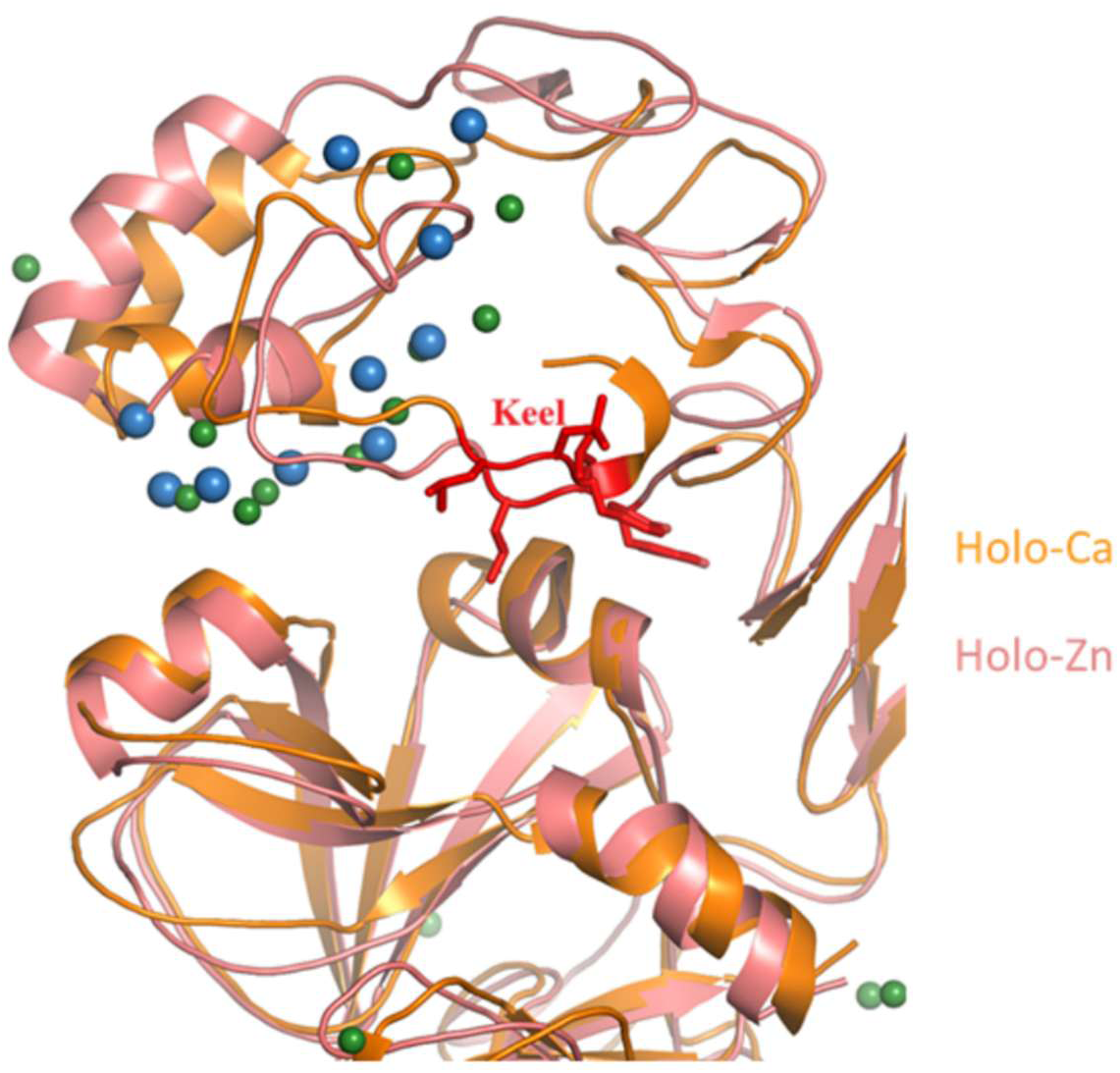
Superimposition of Ca^2+^ bound and Zn^2+^ bound structures of PZ. Zn^2+^ bound structure was obtained by replacing the Ca^2+^ ions (blue spheres) by Zn^2+^ ions (green spheres) in the closed conformation of PZ-Gla domain and subsequently simulating the resulting coordinates to achieve a stable structure. Zn^2+^ occupied the Ca^2+^ binding sites causing minimal perturbation to the overall structure.

Next, Zn^2+^ ions were introduced into the apo protein and subjected to simulation under conditions identical to those used with calcium, to determine whether Zn^2+^ could induce a similar conformational switch in the PZ structure. Interestingly, Zn^2+^ also promoted a slightly folded conformation in PZ; however, it did not achieve the same degree of folding as observed in the presence of Ca^2+^ (Fig. 6). This limitation was attributed to a more extended conformation of the GGSP residues in the EGF domain in presence of Zn^2+^. Consequently, the Gla domain was found to be more flexible in the presence of Zn²⁺ compared to Ca²⁺. The N-terminal EGF residues, GGSP, are conserved and unique to the PZ family. Our observations suggest that conformational changes in PZ are triggered by rotation around these residues, implicating a potential functional role that warrants further investigation. Additionally, one Zn^2+^ ion was found to occupy a site in the EGF domain, coordinated by Glu 83 and Glu 92 (Fig. 7). The ionic interaction between Zn^2+^ and these two Glu residues may prevent the Gla domain from approaching the SP-like domain, resulting in an extended conformation.

**Fig. 6.**
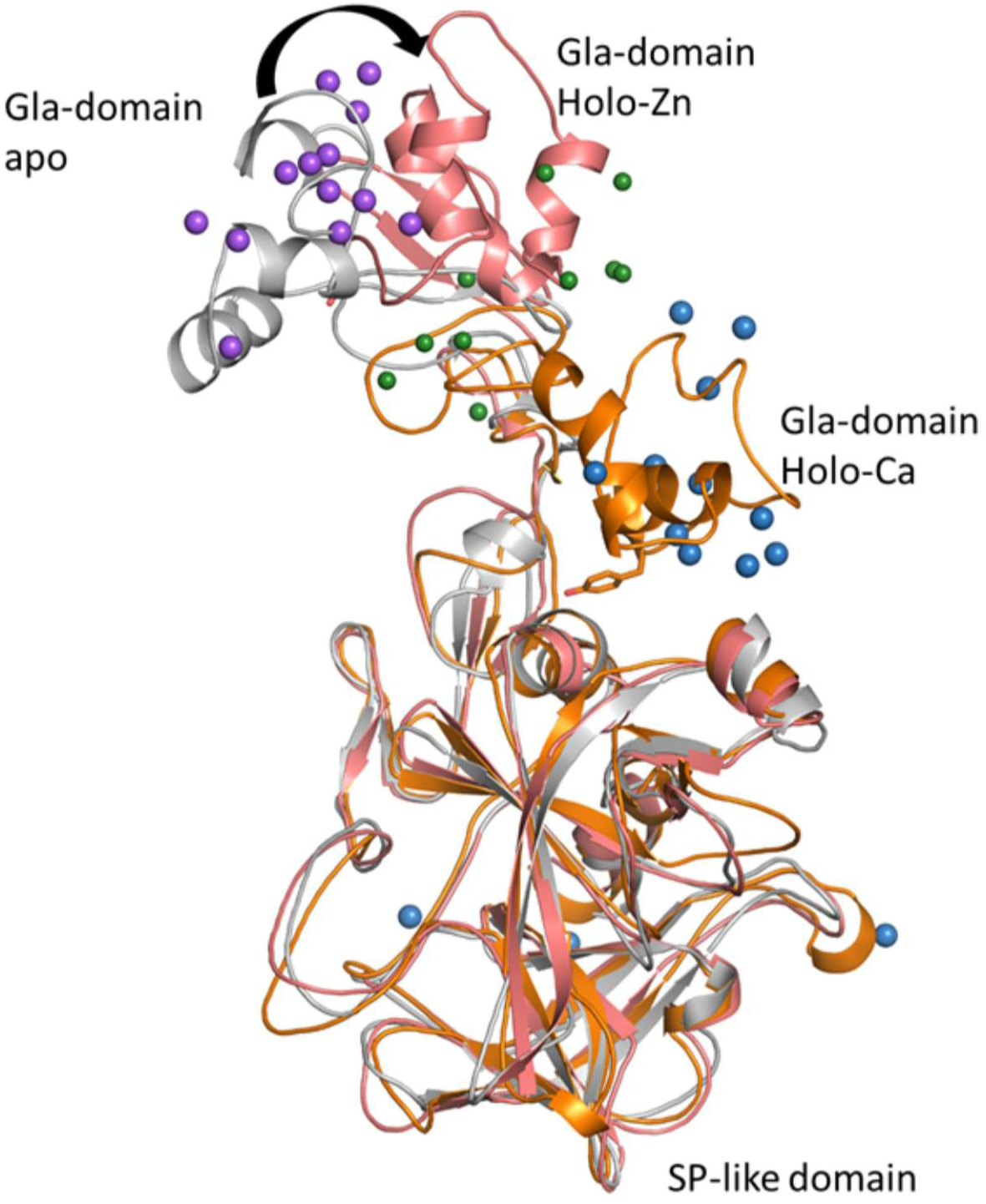
Comparison of holo-Zn^2+^ bound PZ (pink) with apo-PZ (grey) and holo-Ca^2+^ bound PZ (orange) structures. Zn^2+^ bound structure was derived by introducing Zn²⁺ ions into the Gla domain of the apo protein, followed by molecular dynamic simulations performed under the same conditions applied to generate Ca^2+^ bound structure.

**Fig. 7.**
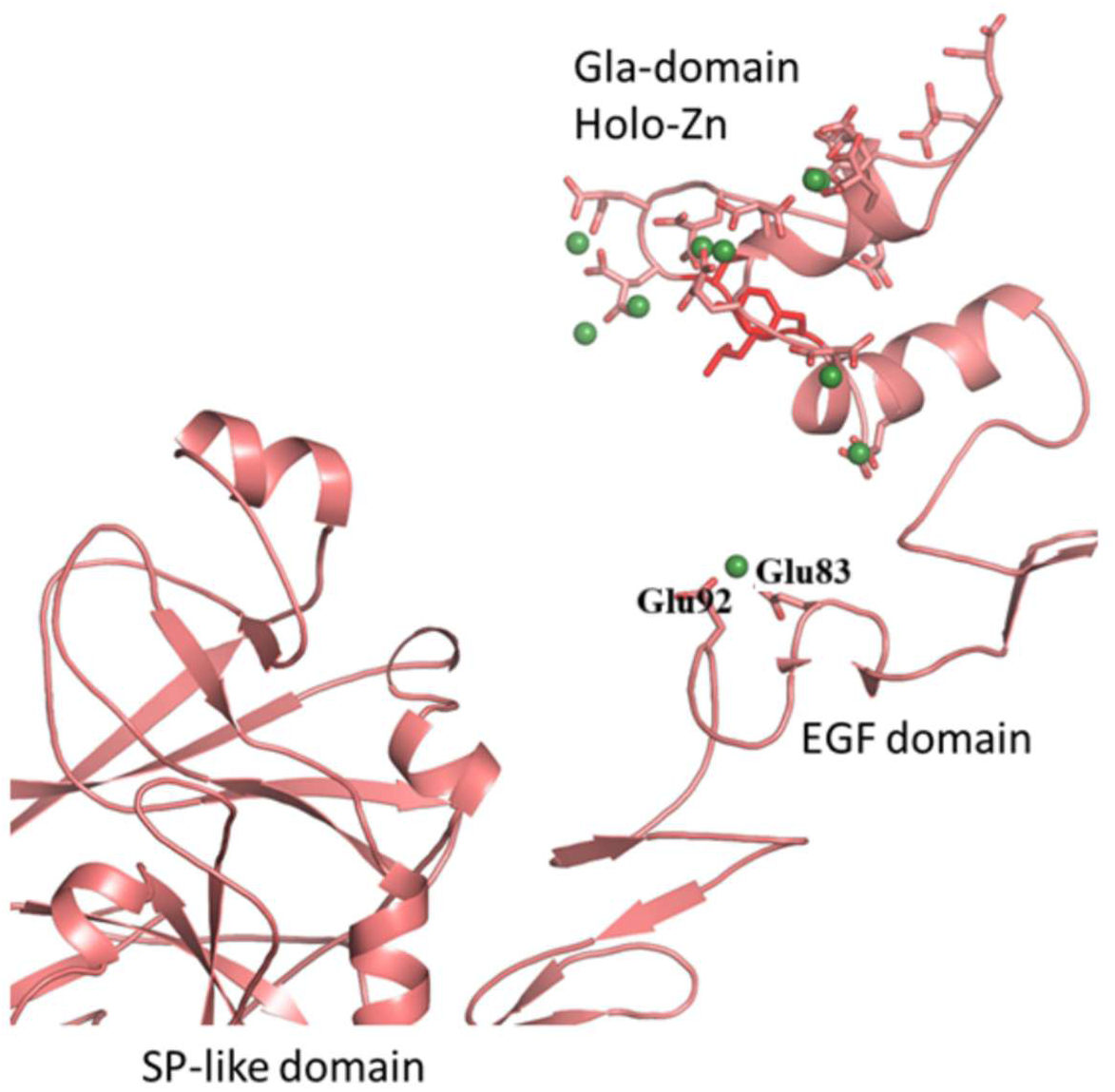
Presence of one Zn^2+^ in EGF domain can impact the domain conformation and inter-domain proximity. Interaction of Zn²⁺ with the carboxylate side chains of Glu83 and Glu92, might promote a stabilized tertiary arrangement that prevented Zn bound Gla domain to approach the SP-like domain via EGF domain rotation.

Finally, we investigated whether the ionic interaction, prompted by Zn^2+^ in the EGF domain, could also restrict Ca^2+^ bound PZ from adopting the folded conformation typically observed. Our MD simulation data revealed that the overall PZ structure retained the same folded conformation as in the original Ca^2+^ bound form, however, the keel region became exposed to the solvent (Fig. 8). Reorientation of the keel region could enhance membrane anchoring of PZ and might also affect the PZ-ZPI interaction, thereby influencing RCL dynamics for fXa inactivation. Thus, Zn^2+^ in the EGF domain of PZ could play a significant role in structural and functional modulation of PZ-ZPI system, acting synergistically with Ca^2+^ in the Gla domain.

**Fig. 8.**
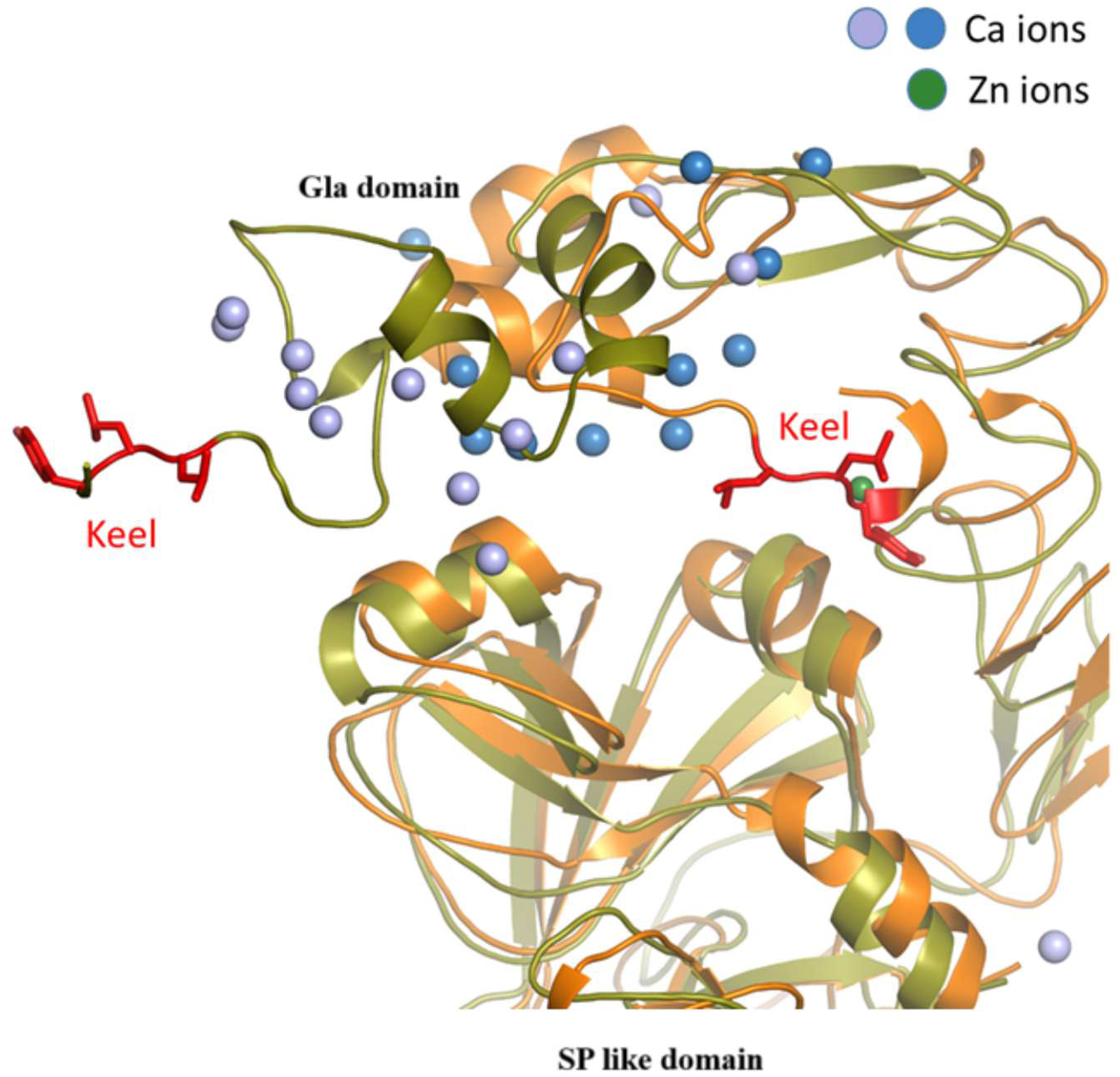
Effect of Zn^2+^ present in EGF domain on Ca^2+^ bound PZ structure. Super imposition of final snapshot of simulated PZ structures in absence (orange) and presence (green) of one Zn^2+^ (green sphere) in EGF domain shows that Zn²⁺-mediated ionic interaction within the EGF domain could not impede the Ca²⁺(light and deep blue spheres) –bound PZ-Gla domain from adopting its characteristic folded conformation. While the Gla domain maintained the canonical folded structure observed in the Ca²⁺-bound state, the keel region became solvent-exposed in presence of Zn^2+^, which may have greater functional significance.

## 4. Concluding remarks

Gla domain plays a critical role in the membrane binding of several serine proteases of coagulation cascade, operating in a Ca^2+^ dependent manner. Each Gla residue carries a –2 charge and coordinates with Ca^2+^ or other bivalent metal ions to facilitate binding to the negatively charged lipids of the membrane. This lipid binding, mediated by the Gla domain, is essential for the coordinated function of the hemostatic system. Concurrently, alterations in membrane binding affinity, whether arising from natural or synthetic variants of VKD coagulation proteins, can significantly impact their coagulant or anticoagulant activities. For example, two missense mutations in the Gla domain of protein S were identified in a patient with protein S deficiency and recurrent thrombotic events. In vitro recombinant studies demonstrated that these mutations impaired the protein’s ability to bind to the membrane, resulting in a functionally compromised variant of protein S [32]. Conversely, selective mutagenesis of the Gla domain in protein C enhanced both its anticoagulant activity and membrane affinity [33]. A chimeric protein C containing prothrombin Gla domain exhibited altered membrane lipid specificity and enhanced anticoagulant activity [34]. Additionally, a Gla domain variant of fVIIa with increased membrane affinity demonstrated augmented procoagulant function [35]. Interestingly, the conserved phosphatidyl serine (PS) binding motif present within the Gla domain of VKD proteins can be used as a molecular probe for imaging, diagnosis, or therapeutic applications targeting cells that externalise PS [36]. Furthermore, a fluorophore conjugated, isolated Gla domain of protein S effectively recognized and bound to PS exposed on diverse cellular surfaces. Collectively, the Gla domains of coagulation proteins serve a crucial role in ensuring proper hemostatic function. PZ, although not a classical serine protease, associates with cellular membranes via its Gla domain and functions as a cofactor for ZPI, facilitating the inhibition of both membrane bound fXa as well as prothrombinase bound fXa during prothrombin activation [37]. Despite this functional importance, the structural mechanism by which metal ions prime the Gla domain for membrane interaction remained unclear, primarily due to the lack of a solution structure for the PZ-Gla domain. The present study addresses this gap by providing comprehensive insights into the structural rearrangements of the PZ domain structure in the presence of divalent metal ions such as Ca²⁺ and Zn²⁺, which are essential for membrane binding. Both Ca²⁺ and Zn²⁺ not only induce the conformational orientation required for effective membrane association and subsequent co factor activity of PZ, but also has a significant contribution in PZ-ZPI interaction and complex formation. Nevertheless, further biochemical characterization is warranted to substantiate these findings. Recent report by Huang et al have demonstrated that PZ independent function of ZPI (D293A ZPI) significantly inhibits occlusive thrombosis while exerting minimal impact on hemostasis [38]. Plasma PZ level was also shown as an independent risk factor of acute ischemic stroke [39]. The insights into membrane-PZ-ZPI interactions gained from our study may offer multiple avenues for designing therapeutic strategies targeting those pathological conditions associated with the PZ-ZPI system.

## CRediT authorship contribution statement

**Subash Chellam Gayathri:** Writing-original draft, review and editing, methodology, investigation, formal analysis. **Suchetana Gupta:** investigation, methodology, formal analysis. **Tanusree Sengupta:** Writing-original draft, review and editing, conceptualization, supervision, funding acquisition. **Soumyadev Sarkar**: methodology, formal analysis

## Acknowledgement

This work was supported by Core Research Grant (CRG/2021/002239) of Science and Engineering Research Board (SERB), Govt. of India to T.S. We gratefully acknowledge the support of the Google Cloud Research Credits (GCRC) program and the high-performance computing (HPC) resources provided by Fluid Numerics to T.S. and S.G. We also thank Prof. A. Gopala Krishna, Department of Biotechnology, Indian Institute of Technology Madras, for access to instrument facility.

