## Supplemental for "Structural Insights into Zn^2+^ and Ca^2+^ binding to Human Protein Z and Their Impact on Membrane Association"

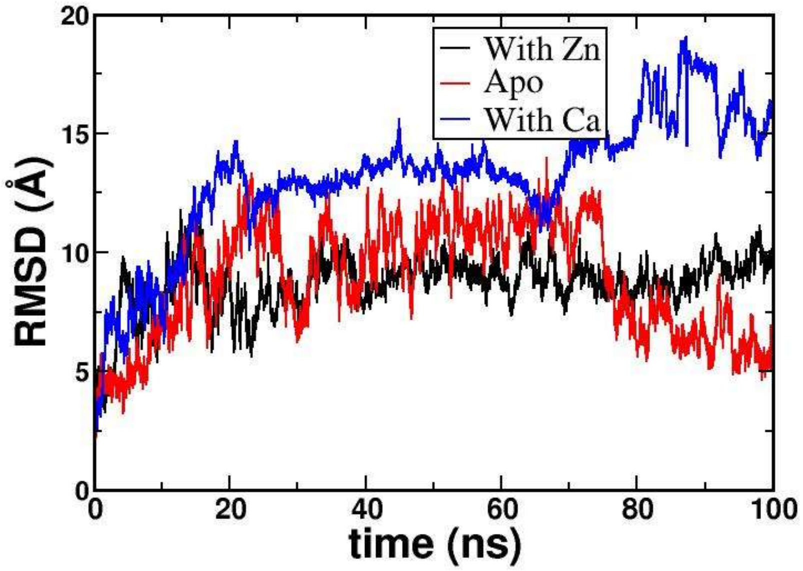


**Fig. S1**. **Plot of backbone root mean square deviation (RMSD) of PZ over 100 ns MD-simulation under three different conditions.** The time evolution for RMSD of the simulated structures with respect to the first frame of simulation has been shown for apo-PZ (red), $Ca^{2+}$ bound PZ (blue), and $Zn^{2+}$ bound PZ (black). Each plot is an average of three simulations.

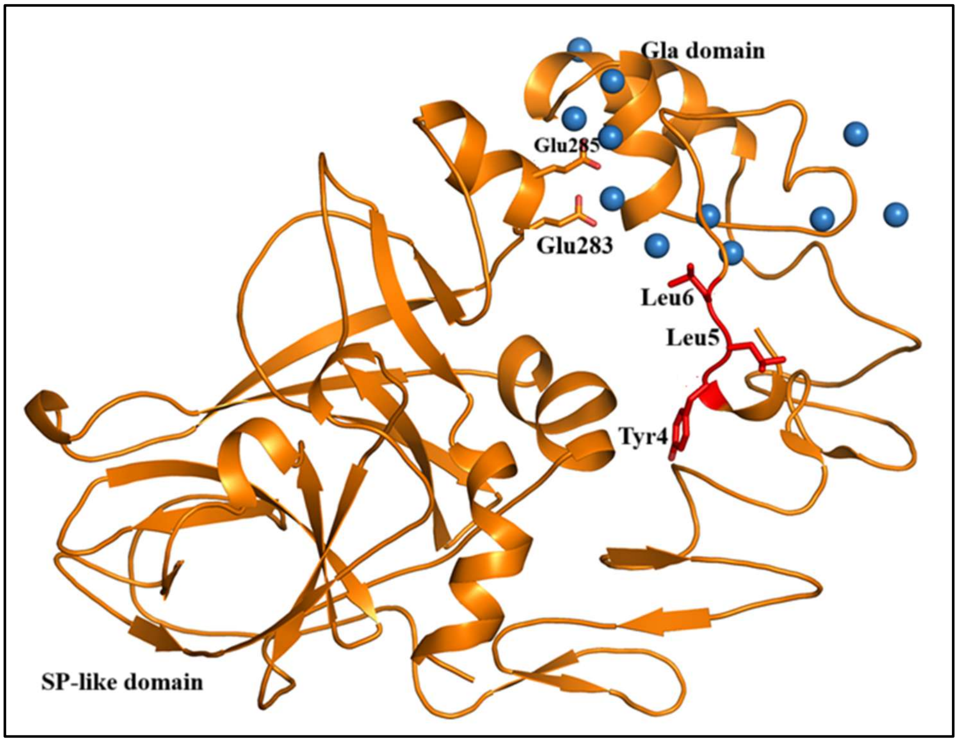


**Fig. S2. $Ca^{2+}$ bound folded PZ structure showing important residues stabilizing folded conformation of PZ.**
